# Analysis of a Novel Bioreactor Designed for Ultrasound Stimulation of Cell-Seeded Scaffolds

**DOI:** 10.1101/476762

**Authors:** Jacob Crapps, Abdolrasol Rahimi, Natasha Case

## Abstract

Although daily low intensity pulsed ultrasound (LIPUS) treatment has been shown to induce cellular responses supporting bone repair, *in vitro* studies in 3D models, such as cell-seeded scaffolds, are needed to further investigate the underlying cellular mechanisms. This requires well-controlled conditions in an US bioreactor. Computational studies are needed to investigate various effects on US wave propagation influenced by bioreactor configurations, such as reflections at interfaces and wave interference, and optimize the bioreactor design for experimental repeatability. In this study, an enclosed cylindrical sample holder that contained an inner well for placement of a scaffold immersed in culture medium was fabricated by stereolithography 3D printing and combined with an acoustic absorbent material to eliminate the presence of an air-liquid interface perpendicular to the wave propagation path. Finite element simulations conducted in the frequency domain demonstrated that weak standing waves were present within the culture medium, indicating the effects of reflections at solid-liquid interfaces within the sample holder, as expected. Focusing on the acoustic pressure at the inner well surface, it was found that the spatially-averaged pressure varied from a maximum to a minimum value as the thickness of the water layer beneath the sample holder was changed. Average pressure values at antinode positions were 2-fold higher than at node positions. A volume-averaged pressure was calculated within the culture medium corresponding to the region where a scaffold would be centrally located within the bioreactor. It was shown that the thickness of the volume analyzed had a minimal effect on the calculated average pressure. Time-dependent simulations for one complete pulse (i.e. 1 ms) showed that the acoustic pressure in volumes that would be occupied by scaffolds of two different thicknesses (diameter of 8.5 mm and thicknesses of 0.2 or 2.0 mm) reached a stable value after 45 µs, and then remained at that value until the active period of the pulse ceased. Once the active period ended, the acoustic pressure rapidly decreased to a low baseline pressure. Overall, this study showed that the proposed novel bioreactor design provided a controlled environment for the US treatment of a cell-seeded scaffold by removing the air-liquid interface using a custom-designed sample holder and an acoustic absorbent material.

## I Introduction

Daily application of a low intensity pulsed ultrasound (LIPUS) regimen has been shown to have positive effects on bone fracture healing [1]. The standard LIPUS protocol applies sinusoidal acoustic waves at 1.5 MHz frequency with a pulsed signal (20% duty cycle and 1 KHz pulse repetition frequency) delivered as 200 µs active time following by 800 µs inactive time for 20 minutes at a spatial and temporal average intensity of 30 mW/cm^2^ [2]. Although LIPUS stimulation can induce cellular responses supporting bone repair [3], the underlying mechanisms are not well understood. While most *in vitro* studies have focused on US stimulation of two-dimensional (2D) cell monolayers, a fracture site requires treatment of a three-dimensional (3D) volume and US propagation within such a space is complex. Therefore, *in vitro* studies in 3D models, such as cell-seeded scaffolds, are needed to investigate the bioeffects of the LIPUS protocol under well-controlled conditions.

*In vitro* experiments have been conducted in various US bioreactor designs [4–7]. A major limitation present in the majority of commonly used bioreactor designs is the presence of an air-liquid interface in the direct path of US wave propagation. When an air-liquid interface is present in the bioreactor, complete wave reflection will occur, resulting in the formation of standing waves [8]. This effect increases variability in the pressure delivered to the cell layer, thereby reducing reproducibility across experiments and the reliability of the US bioreactor. To provide a well-controlled environment for ultrasound stimulation of cell-seeded scaffolds, an ideal bioreactor configuration would a) eliminate air-liquid interfaces along the US wave transmission path; b) allow for the central positioning of the cell-seeded scaffold; c) support quick assembly and disassembly of the bioreactor for daily use in a multi-day stimulation regimen; and d) provide a way to trap any air bubbles introduced into the culture medium during bioreactor assembly away from the central zone of the bioreactor.

Computational modeling is a valuable approach to study US wave propagation in the bioreactor and allows researchers to study the acoustic pressure field in regions where experimental measurement is not possible. For instance, pressure cannot be measured using a hydrophone at the dish surface where the cell layer is present in a 2D culture model or within a porous scaffold where cells will be distributed in a 3D culture model. A few previous studies have examined variability of the pressure field in the US bioreactor and evaluated effects of geometric features [9, 10]. One limitation of these prior studies was that the acoustic pressure field was simulated only in the frequency domain (which assumes continuous waves) or only for a short period of time (over a few sine waves).

The purpose of this study was to develop and characterize a novel bioreactor configuration for US stimulation of cell-seeded scaffolds that eliminated all air-fluid interfaces perpendicular to the wave propagation path. In our bioreactor configuration, an enclosed sample holder filled with culture medium, was combined with an acoustic absorbent material. The US transducer was immersed in water and positioned below the sample holder. The acoustic pressure in the fluid domains of the US bioreactor was evaluated by developing a finite element model and conducting multiphysics simulations. The effects of individual configuration parameters (i.e. thicknesses of the water and culture medium layers) on the volume-averaged acoustic pressure over a region that a cell seeded scaffold would occupy were analyzed in the frequency domain. In a subset of simulations, time-dependent pressure variations generated by LIPUS were analyzed by simulating the acoustic pressure field over one pulse (i.e. 1 ms).

## II METHODS

### A Development of a Sample Holder

A novel sample holder was designed and fabricated. The 3D CAD model of the sample holder was designed in Autodesk Inventor (San Rafael, CA) and printed using a stereolithography technique (Form 2, Formlabs Inc., Somerville, MA) with a clear resin (ρ = 1170 kg/m3 and E=2.8 GPa). As shown in Figure 1, two pieces were designed to screw together creating a central horizontal channel filled with culture medium through which the US waves would pass and an outer vertical cylindrical channel that provided a place for trapping air bubbles away from the US wave transmission path. The base of the holder had a central well into which a scaffold could be placed to position it at the center of the US pathway.

**Figure 1.**
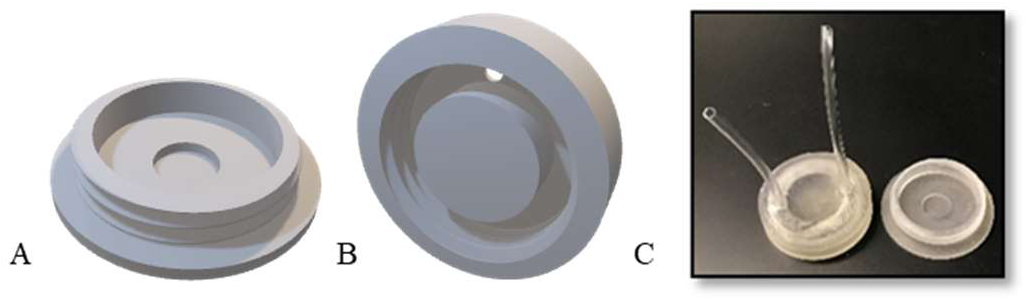
(A) CAD model of the base piece of the sample holder with the inner chamber used to position a cell-seeded scaffold. (B) CAD model of the top piece of the sample holder containing an outer channel to capture air bubbles. (C) 3D printed sample holder using clear resin, with plastic tubing attached to the top piece to provide a route for gas exchange and the ability to fill the holder with culture medium.

In order to estimate Poisson’s ratio for the clear resin, it was necessary to experimentally determine the speed of sound in the resin. The sample holder base was filled with water and a needle hydrophone (400 μm active element diameter; HNP-400, Onda Corp., Sunnyvale, CA) connected to a pre-amplifier (AH-2010, Onda Corp.) was positioned in the water. The sample holder was placed on the surface of a water tank, with a US transducer with a center frequency of 1.5 MHz (TXCP, Precision Acoustics Ltd, Dorchester, UK) immersed in the water under the sample holder.

Using a function generator (DG1022, Rigol Technologies, Beijing, China) connected to the US transducer, a short burst of waves (10 sine waves) were sent and the arrival time was measured. The hydrophone output was recorded via an oscilloscope (DSOX3024T InfiniiVision, Keysight Inc., SantaRosa, CA). The speed of sound in the clear resin was calculated as follows [11]:

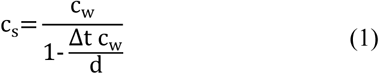

Here, c_s_ and c_w_ are the speed of sound in the clear resin and in water, respectively. Δt is the average change in arrival time of the US waves when the sample holder is placed between the US transducer and the hydrophone, and *d* is the thickness of the sample holder bottom thickness.

The measurement was repeated five times to generate an average value for *C*_s_, which was implemented in the model. The density of the clear resin was calculated simply by measuring the weight and volume of a 3D printed solid block of this material.

### B. Finite Element Modeling

A finite element model of the US bioreactor (Figure 2) was developed in COMSOL Multiphysics software (V5.3, COMSOL Inc., Stockholm, Sweden). The US transducer model contained a piezoelectric disk, a backing layer, a matching layer, and a stainless steel casing. The backing layer, made of epoxy/tungsten, absorbs US waves propagating from the back of the piezoelectric disk and supports the disk. The matching layer, made of epoxy, reduces the acoustic impedance mismatch between the piezoelectric material and the water, so that maximum energy is transferred out of the transducer. An acoustic absorbent material (Aptflex F28, Precision Acoustics, Dorchester, UK) was placed on top of the sample holder, and also surrounded the water layer between the transducer and sample holder. At a frequency of 1.5 MHz, this material absorbs over 99% of incoming US waves while reflecting very little. Having this material above and around the bioreactor effectively eliminates air-liquid boundaries where US reflections can occur. The material properties for the components of the model are listed in Table 1. The pressure wave speeds used for culture medium, water, and air were 1515, 1471, and 344 m/s, respectively.

**Table 1.** The physical properties and geometrical parameters of materials implemented in the finite element model. E:elastic modulus.

| Material | Density [Kg/m<sup>3</sup>] | E [GPa] | Poisson's Ratio | Stiffness Damping Parameter | Thickness [mm] |
| --- | --- | --- | --- | --- | --- |
| Epoxy | 1096 | 5.5 | 0.34 | $2.3 \times 10^{-8}$ | 0.375 |
| Piezoelectric | 7500 | - | - | $5.3 \times 10^{-10}$ | 0.75 |
| Epoxy/Tungsten | 5766 | 10.5 | 0.34 | $1.5 \times 10^{-8}$ | 30.6 |
| Stainless Steel | 7900 | 193 | 0.29 | $1 \times 10^{-7}$ | 3.2 |
| Clear resin | 1170 | 2.8 | 0.45 | $4 \times 10^{-9}$ | 0.3<br>(inner well) |
| Culture Medium | 1004 | - | - | - | - |
| Water | 998 | - | - | - | - |
| Air | 1.24 | - | - | - | - |
| Acoustic Absorbent | 1010 | 0.259 | 0.48 | 10 | 10 |

**Figure 2.**
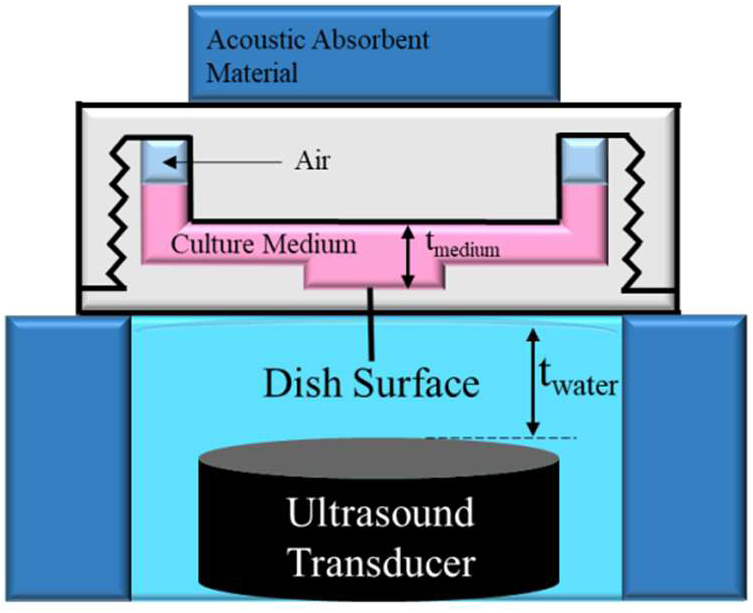
A computational model was created for ultrasonication from below the dish. The acoustic pressure was evaluated as a spatial average of the culture medium layer contacting the surface of the inner well or as a volume average of a centrally located cylindrical region of the culture medium that corresponded to the volume of a scaffold of radius 4.5 mm and of specified thickness.

### C. Governing Equations

To analyze US wave propagation in the sample holder and through the fluid and solid layers, three physics were implemented in the model. The electric displacement field (D) and the electric potential (E) in the piezoelectric disk were simulated with the constitutive equation as below,

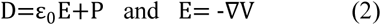

Here, ɛ_0_ [F/m] is permittivity of the vacuum, P [C/m^2^] is the electric polarization vector, and ∇V [V] is the applied electric potential gradient.

A solid mechanics interface was used to apply an isotropic, linear elastic material property to the culture dish and the transducer components. The constitutive equation was defined by Hooke’s law:

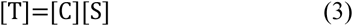

Here, T [Pa] is the stress tensor, C [Pa] is the elastic coefficients, and S [m‧m^-1^] is the strain tensor.

The acoustic pressure in the fluid domain was simulated in the frequency domain using the Helmholtz wave equation,

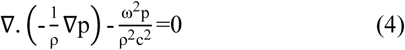

And for the time-dependent model as shown:

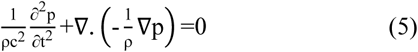

Here, ρ [kg/m^3^] is density, p(z,t)=p(z).e^iwt^ [Pa] is acoustic pressure, ω [rad/s] is the angular frequency, and c [m/s] is the pressure wave speed.

The piezoelectric property was defined by the constitutive equations:

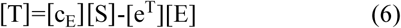

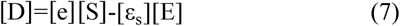

Here, c_E_ [Pa] is the matrix of elastic coefficients at the constant electric field, e^T^ [C/m^2^] is the transposed coupling matrix, e [C/m2] is the coupling matrix, and ɛ_s_ [F/m] is the permittivity matrix at the constant mechanical strain.

#### D. Boundary Conditions and Attenuation

The lower surface of the piezoelectric disk was grounded (Voltage=0 V) and a pulsed electric potential of 20V peak-to-peak at 1.5 MHz was applied to the top surface. The pulsed electric potential duration was 1 ms with 10%, 20%, or 50% duty cycle. An Acoustic-Structure Boundary Coupling was added to couple the fluid and solid physics together.

To account for acoustic attenuation by certain components, the common attenuation model of Rayleigh damping was used, with the following constitutive equation:

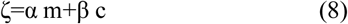

Here, ζ is the damping parameter, α is the mass damping parameter, β is the stiffness damping parameter, m [kg] is the mass, and c [N/m] is the stiffness. Attenuation in the water and culture medium were considered negligible and not included in the model. The mass damping parameter was set to zero in all materials in which attenuation was included. The stiffness damping parameters used in the model are listed in Table 1.

#### E. Mesh / Study Type

A triangular mesh type for both a time-dependent solver and a frequency-dependent solver were used. The mesh element size and time step were calculated by equations as below:

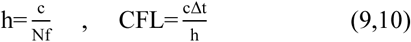

Here, h [mm] is the mesh element size, f [Hz] is the frequency, and c [m/s] is the speed of sound. N, the number of mesh elements per wavelength, was chosen as 8 after conducting a mesh convergence analysis. The CFL (Courant-Friedrichs-Lewy) number was set to the optimal value of 0.2 to calculate Δ*t* [s], the time step used in the time-dependent solver, and Δ*t*=13 ns was used in the solver. The mesh element size was different in each material and depended upon the sound wavelength.

## III. RESULTS AND DISCUSSION

This study focused on using computational analyses to evaluate how variations in individual bioreactor configuration parameters (i.e. thicknesses of the water and culture medium layers) changed the absolute acoustic pressure in the culture medium layer within the central well of the sample holder. With the transducer positioned below the culture dish and immersed in water, US waves traveled through each consecutive layer of the model including the water, base of the sample holder, culture medium, top portion of the sample holder, and the acoustic absorbent material (Figure 2).

The solid-liquid interfaces present in this configuration produced partial wave reflection (~34% at each interface), generating repeating layers of maximum and minimum acoustic pressures along the z-axis within both the culture medium and water layers (Figure 3). The regions of maximum acoustic pressure resulted from maximum constructive interference of transmitted and reflected waves and will be referred to as antinode positions, while the regions of minimum acoustic pressure resulted from maximum destructive interference of transmitted and reflected waves and will be referred to as node positions. Comparing the pressure field at an antinode position (Figure 3.A) and at a node position (Figure 3.B) showed limited change in the overall pressure pattern, but the maximum pressure level experienced at the node position was 48% lower.

**Figure 3.**
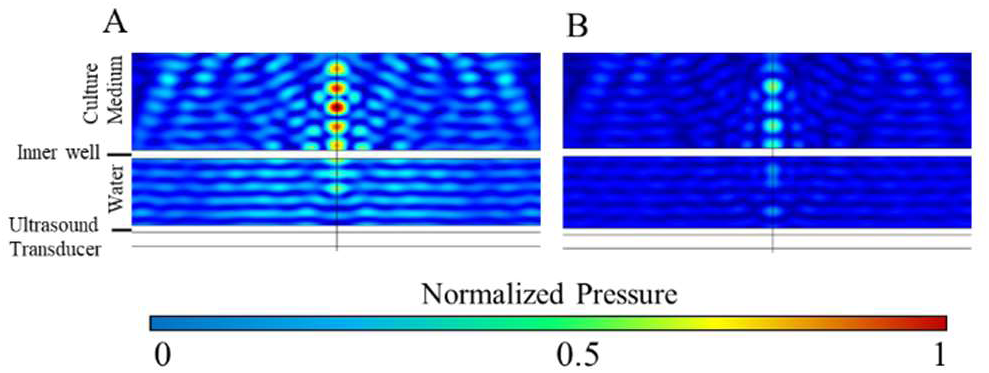
Comparison of the acoustic pressure field in the fluid layers of the bioreactor for two different water layer thicknesses for a frequency domain simulation. (A) Antinode location (t_water_=2.44 mm) where maximum constructive wave interference occurred at the interface of the culture medium and well surface. (B) Node location (t_water_=2.20 mm) where maximum destructive wave interference occurred at the interface of the culture medium and well surface. All results were normalized to the maximum pressure experienced within the culture medium for the antinode location and t_medium_=3.6 mm.

To better understand the influence of the water layer thickness on acoustic pressure at the well-culture medium interface, simulations in the frequency domain were run in which the thickness of the water layer was varied over a 2 mm span using a step size of 50 µm. Simulations were conducted for four different distances between the sample holder base and transducer: 2 – 4 mm, 24 – 26 mm, 49 – 51 mm, and 74 – 76 mm. Overall, these simulations showed that the spatially-averaged acoustic pressure at the well surface varied from a maximum to a minimum value as the thickness of the water layer beneath the sample holder was changed by approximately 0.245 mm, which is equal to *λ_medium_*/4 (Figure 4.A – 4.D).

**Figure 4.**
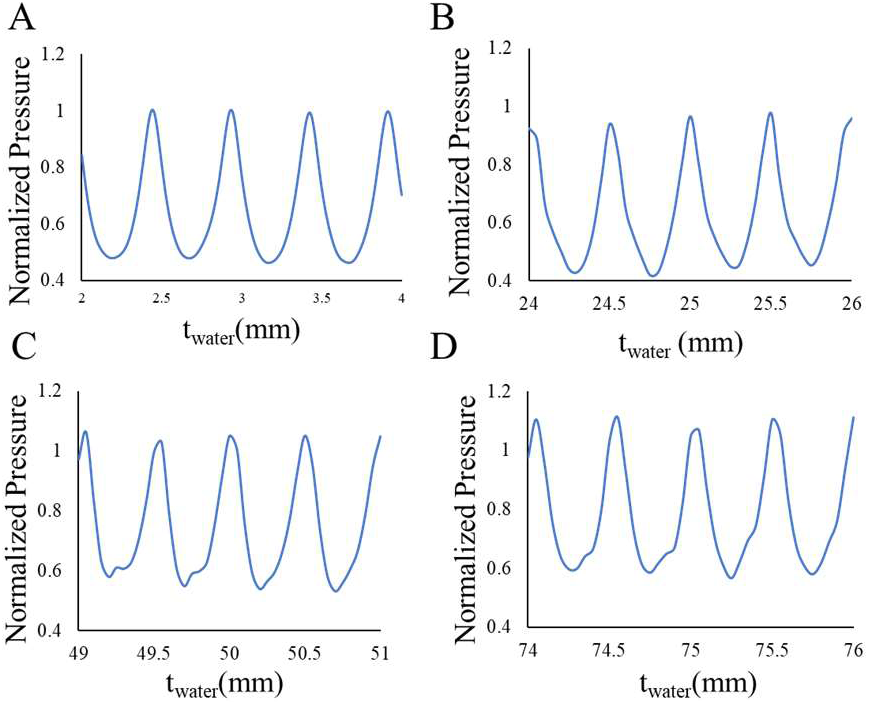
Evaluating the sensitivity of the spatially-averaged acoustic pressure at the inner well surface to the change of the water layer thickness at different regions. All results are normalized to the average pressure at antinode locations in the 2-4 mm water thickness range (t_medium_=3.6 mm).

The acoustic pressure at the well surface for the antinode locations was 2.1 times higher than at the node locations for the four consecutive node and antinode positions shown in Figure 4.A. Similarly, the ratio of the acoustic pressure at the well surface of the antinode to node positions for the regions of 24 mm < t_water_ < 26 mm, 49 mm < t_water_ < 51 mm, and 74 mm < t_water_ < 76 mm were 2.2, 1.9, and 1.7-fold, respectively. The results showed that the sensitivity of the acoustic pressure at the well surface to the water layer thickness was approximately similar at all four regions evaluated. For the graph shown in each subfigure, the pressure was averaged and the average values in the ranges of 24 mm < t_water_ < 26 mm, 49 mm < t_water_ <51 mm, and 74 mm < t_water_ < 76 mm were 97%, 122%, and 134% of the average value at the range of 2 mm < t_water_ < 4 mm.

To analyze the effect of the culture medium layer thickness on acoustic pressure at the well-culture medium interface, simulations were run for t_medium_ changing from 3.2 to 5.2 mm, with the thickness of the water layer set to an antinode location of t_water_=2.44 mm. Figure 5.A shows that changing the culture medium layer thickness by approximately 0.253 mm, which is equal to *λ_medium_*/4, shifted the acoustic pressure at the well surface between a node and antinode position. On average, the acoustic pressure dropped by 36% when the culture medium was either increased or decreased to shift the well surface from an antinode to a node position. Note that t_medium_=3.6 mm was used for all other simulations. Using these results, a sample holder can be fabricated to have the desired culture medium height corresponding to an antinode position for a set water layer thickness. The effect of culture medium height variation was previously investigated for a traditional ultrasonication from below configuration and it was reported that an 18 µL variation in 13 mL culture medium in a well can lead to a 10% variation in the pressure at the dish surface [9]. The design of our sample holder allows the same culture medium height to be used between experiments, thereby removing this configuration parameter as a source of variability and enhancing the reproducibility of our experimental results.

**Figure 5.**
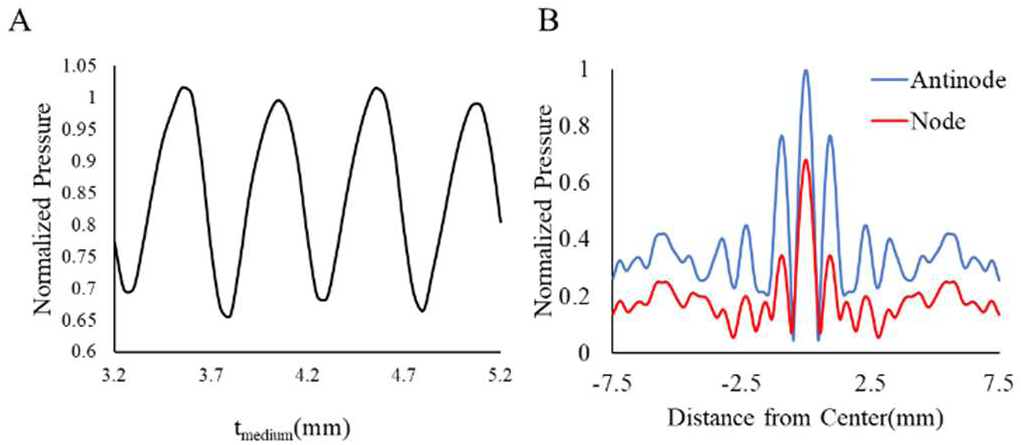
Analyzing the effects of the culture medium layer thickness on the acoustic pressure at the well surface. (A) Spatially averaged pressures varied with the thickness of the culture media layer. (B) The radial acoustic pressure pattern at the well surface at antinode (t_medium_= 3.6 mm) and node (t_medium_= 3.8 mm) locations. Results are normalized to the average antinode pressure in (A) (t_water_= 2.44 mm).

The radial pressure pattern at the well surface was investigated, as shown in Figure 5.B. The maximum pressure occurred at the center and dropped by 74% at the distance of 7.5 mm from the center when located at an antinode position. The pressure drop under the same condition was 81% when located at a node position. Overall, the pattern in acoustic pressure variation was approximately similar for both node and antinode positions, with the pressure being lower at the node position for the majority of radial positions.

Fig. 5B, along with Fig.3, illustrated a potential design flaw in the sample holder. The radial pressure values were approximately 60% of the maximum value in the center of the chamber when located 2 mm or farther from the center. Also, as shown in Fig. 3, the layers of local maxima and minima were not uniform. This could result in a highly nonuniform pressure field for stimulation of cells within a scaffold, with cells in the center receiving a much higher pressure than cells at the edge of the scaffold.

A simulation was conducted to compare the effect that the water layer thickness had on volumetric averages representing the region where a cell-seeded scaffold would be positioned within the sample holder, with the thickness of the water layer varied from 2-4 mm (Fig.6A & B). The average acoustic pressure was analyzed for a cylindrical volume with a a radius of 4.25 mm and a thickness of either 0.2 mm or 2.0 mm, which were chosen to represent volumes of thin and thick scaffolds, respectively. The pressure in the 2mm thick volume was on average 84% of the pressure in the 0.2 mm thick volume. Antinode pressures were 17% greater when analyzed over a thickness of 0.2 mm when compared to a thickness of 2 mm (Fig. 6A). To further analyze the effect of cylindrical volume thickness on the volume-averaged pressure in the culture medium, a simulation was conducted when both the culture medium and water layer thicknesses were set to antinode positions (t_water_= 2.4 mm and t_medium_=3.6 mm; Fig. 6B). Volume-averaged acoustic pressures were calculated as before. Before the thickness exceeded 0.5 mm, the maximum change in volume-averaged pressure was 14%. After the thickness of the volume analyzed exceeded 0.5 mm, there was minimal variation in the calculated volume average (≤ 2%).

**Figure 6.**
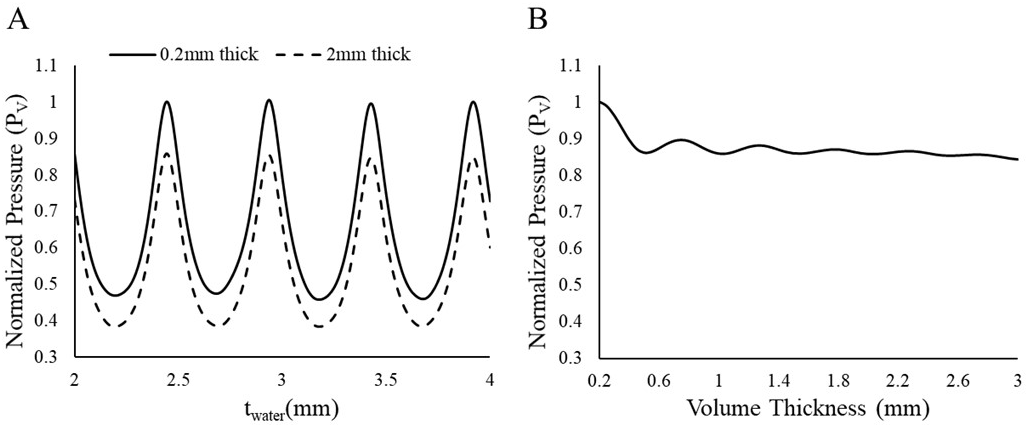
Average pressures were analyzed throughout volumes of different thicknesses (A) The thickness of the water layer was varied from 2-4 mm, and the average pressure in two different volumes was normalized to the average antinode pressure over the 0.2 mm thick volume. (B) The volume of analysis for average acoustic pressure was varied with thicknesses from 0.2 to 3 mm, with a set radius of 4.25 mm. The values were normalized to the same value as in (A). P_V_ denotes volume averaged pressure as the values being normalized.

Time-dependent simulations were conducted over one pulse (1ms) for the standard pulsed US protocol (20% duty cycle). This simulation was conducted in order to better illustrate the varying acoustic pressure that would occur within the culture medium during one US pulse. The standard protocol was chosen due to the complexity and time required to compute time-dependent simulations at a frequency of 1.5 MHz. The two cylindrical volumes representing thin and thick scaffold dimensions, as evaluated in Figure 6, were analyzed, and Figure 7 shows the average sinusoidal acoustic pressure with respect to time. With initiation of the US signal, the acoustic pressure increased until it reached a stable value, and then did not change for the rest of the active period. The time to reach to the stable pressure value was 45 µs for both volume regions. As the US signal ceased, the average acoustic pressure dropped rapidly to a low baseline pressure. The stable acoustic pressure value was 11% lower when the pressure was averaged over a volume corresponding to a 2 mm thickness compared to a volume corresponding to a 0.2 mm thickness.

**Figure 7.**
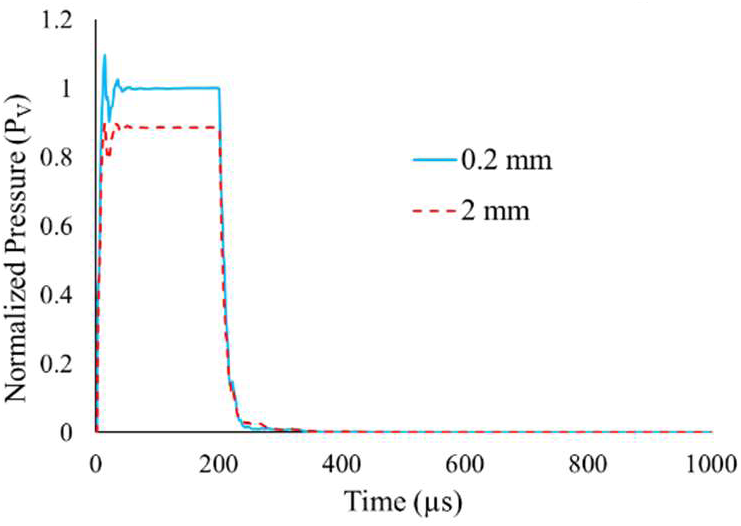
Using a time-dependent simulation, the acoustic pressure pattern over 1ms was analyzed. Pressures were averaged over the volume regions with diameter of 8.5 mm and thicknesses of 0.2 and 2 mm. For the displayed results, each data point represents the time average pressure over one wave period (0.667 µs). Results were normalized to the stable acoustic value of the 0.2 mm thick volume region (t_water_= 2.44 mm, t_medium_=3.6 mm).

## IV. CONCLUSION

Overall, this study showed that the proposed novel bioreactor design provided a controlled environment for the US treatment of a cell-seeded scaffold. It effectively removed all air-liquid interfaces using a uniquely designed sample holder and an acoustic absorbent material. The sample holder also prevented possible scaffold movement during the experiment through the use of an enclosed chamber with an inner well to position cell-seeded scaffolds. However, this study did have limitations. Scaffolds were not included as a solid component of the finite element model. Rather, acoustic pressures were averaged over a volume of culture medium that a scaffold would ideally occupy. This simplified approach neglected how the acoustic properties of the porous scaffolds would affect US wave propagation in the bioreactor. Future work would include modeling the scaffolds as porous solids with appropriate acoustic properties. Additionally, this study showed that the proposed geometry of the inner well of the sample holder caused nonuniform US wave propagation in the region where cells would be located. A future improvement to this proposed design would be to adjust the geometry of the well to allow for more uniform US wave propagation, while still maintaining a method for centering the scaffolds in the holder.

